# Seabird morphology determines operational wind speeds, tolerable maxima and responses to extremes

**DOI:** 10.1101/2022.05.02.490292

**Authors:** Elham Nourani, Kamran Safi, Sophie de Grissac, David J. Anderson, Nik C. Cole, Adam Fell, David Grémillet, Emmanouil Lempidakis, Miriam Lerma, Jennifer L. McKee, Lorien Pichegru, Pascal Provost, Niels C. Rattenborg, Peter G. Ryan, Carlos D. Santos, Stefan Schoombie, Vikash Tatayah, Henri Weimerskirch, Martin Wikelski, Emily L. C. Shepard.

**Affiliations:** Department of Migration, Max Planck Institute of Animal Behavior, Am Obstberg 1, 78315 Radolfzell, Germany; Department of Biology, University of Konstanz, Universitätsstraße 10, 78464 Konstanz, Germany; Diomedea Science – Research & Scientific communication, 819 route de la Jars, 38 950 Quaix-en-Chartreuse, France; France Energies Marines, Antenne Méditerranée, 38 rue Frédéric Jolliot-Curie, 13451 Marseille, France; Department of Biology, Wake Forest University, 1834 Wake Forest Rd, Winston-Salem, NC 27109, USA; Durrell Wildlife Conservation Trust, La Profonde Rue, JE3 5BP Jersey, Channel Islands; Biological and Environmental Sciences, University of Stirling, FK9 4LA Stirling, UK; CEFE, Univ Montpellier, CNRS, EPHE, IRD, 1919 route de Mende, 34293 Montpellier, France; FitzPatrick Institute of African Ornithology, DST-NRF Centre of Excellence, University of Cape Town, Rondebosch, 7701 Cape Town, South Africa; Department of Biosciences, Swansea University, SA1 8PP Swansea, UK; Research and Technology Centre (FTZ), University of Kiel, Hafentörn 1, 25761 Büsum, Germany; Institute for Coastal and Marine Research, Nelson Mandela University, 6031 Gqeberha, South Africa; Ligue pour la Protection des Oiseaux, Réserve Naturelle Nationale des Sept-Iles, 22560 Pleumeur Bodou, France; Avian Sleep Group, Max Planck Institute for Ornithology, Eberhard-Gwinner-Straße, 82319 Starnberg, Germany; Mauritian Wildlife Foundation, Grannum Road, 73418 Vacoas, Mauritius; Núcleo de Teoria e Pesquisa do Comportamento, Universidade Federal do Pará, R. Augusto Corrêa, 01 - Guamá, PA, 66075-110, Belém, Brazil; CESAM - Centro de Estudos do Ambiente e do Mar, Faculdade de Ciências, Universidade de Lisboa, Campo Grande 016, 1749-016 Lisboa, Portugal; Centre for the Advanced Study of Collective Behaviour, University of Konstanz, Universitätsstraße 10, 78464 Konstanz, Germany

**Keywords:** Extreme weather events, storms, flight, wing loading, bio-logging

## Abstract

Storms can cause widespread seabird strandings and wrecking^1,2,3,4,5^, yet little is known about the maximum wind speeds that birds are able to tolerate or the conditions they avoid. We analyzed > 300,000 hours of tracking data from 18 seabird species, including flapping and soaring fliers, to assess how flight morphology affects wind selectivity, both at fine scales (hourly movement steps) and across the breeding season. We found no general preference or avoidance of particular wind speeds within foraging tracks. This suggests seabird flight morphology is adapted to a “wind niche”, with higher wing loading being selected for in windier environments. In support of this, wing loading was positively related to the median wind speeds on the breeding grounds, as well as the maximum wind speeds in which birds flew. Yet globally, the highest wind speeds occur in the tropics (in association with tropical cyclones) where birds are morphologically adapted to low median wind speeds. Tropical species must therefore show behavioral responses to extreme winds, including long-range avoidance of wind speeds that can be twice their operable maxima. In contrast, procellariiformes flew in almost all wind speeds they encountered at a seasonal scale. Despite this, we describe a small number of cases where albatrosses avoided strong winds at close-range, including by flying into the eye of the storm. Extreme winds appear to pose context-dependent risks to seabirds, and more information is needed on the factors that determine the hierarchy of risk, given the impact of global change on storm intensity ^6,7^.

## Results

The windscape largely determines seabirds’ flight costs. Seabirds should therefore adapt their behaviors to reduce these costs in strong wind conditions. Wandering albatrosses, for example, travel in large loops from Crozet Island, taking advantage of regional wind patterns to avoid headwinds^8^. Some seabirds reduce exposure to strong winds by adjusting their flight distance to circumnavigate storms^9^. Even more widespread is the tendency to decrease flight height when flying into a headwind^10,9^. Here we investigated whether flight morphology affects wind selectivity, both at fine scales (hourly movement steps) and across the breeding season. We focused on preference or avoidance of wind speeds, rather than wind direction, as the latter could be influenced by prey distribution, competition, and the need to return to a central place ^11^.

To understand the wind speeds seabirds are able to fly in and those they avoid, we analyzed 1,663 foraging trips from 18 species of seabirds (Table S1), representing 326,960 hours of flight time. We compared the wind speeds each bird encountered at every hourly segment (hereafter a step) along its foraging track with wind speeds at 30 alternative steps that were available to the bird in space and time but were not used. Following this step-selection approach (Figure S1), our dataset had a stratified structure, where each stratum was composed of one observed step and its 30 alternative steps.

We chose not to proceed with the conditional logistic regressions commonly used to estimate step-selection functions ^12^, as this disregards the outliers encountered by the birds in flight. Instead, we used null modeling to compare the used and the maximum available wind speed within each step, and thereby identify the rare wind-avoidance events (STAR Methods).

Birds experienced a wide range of wind speeds overall (Figure 1). However, the variation in wind speeds available to them at any one point in time (calculated as the coefficient of variation for wind speed values within each stratum) was generally low, with 90% of the strata showing variation lower than 11% (Figure S2), and used wind speeds were not significantly different from the strongest speeds available to the birds (Figure S3). Maximum wind speeds were avoided in only nine of the 93,104 strata, involving four species: Atlantic yellow-nosed albatross, Wandering albatross, Sooty albatross, and Red-footed booby (Figure S3). In six of the nine avoidance cases, birds avoided wind speeds that were within their population-specific flyable range of wind speeds (Figure 1). The trips containing avoidance behavior indicated that the Atlantic yellow-nosed and Wandering albatrosses were responding to storms by selecting the region of lower wind speeds by flying towards and tracking the eye of the storm (Figure 2; Video S1).

**Figure 1:**
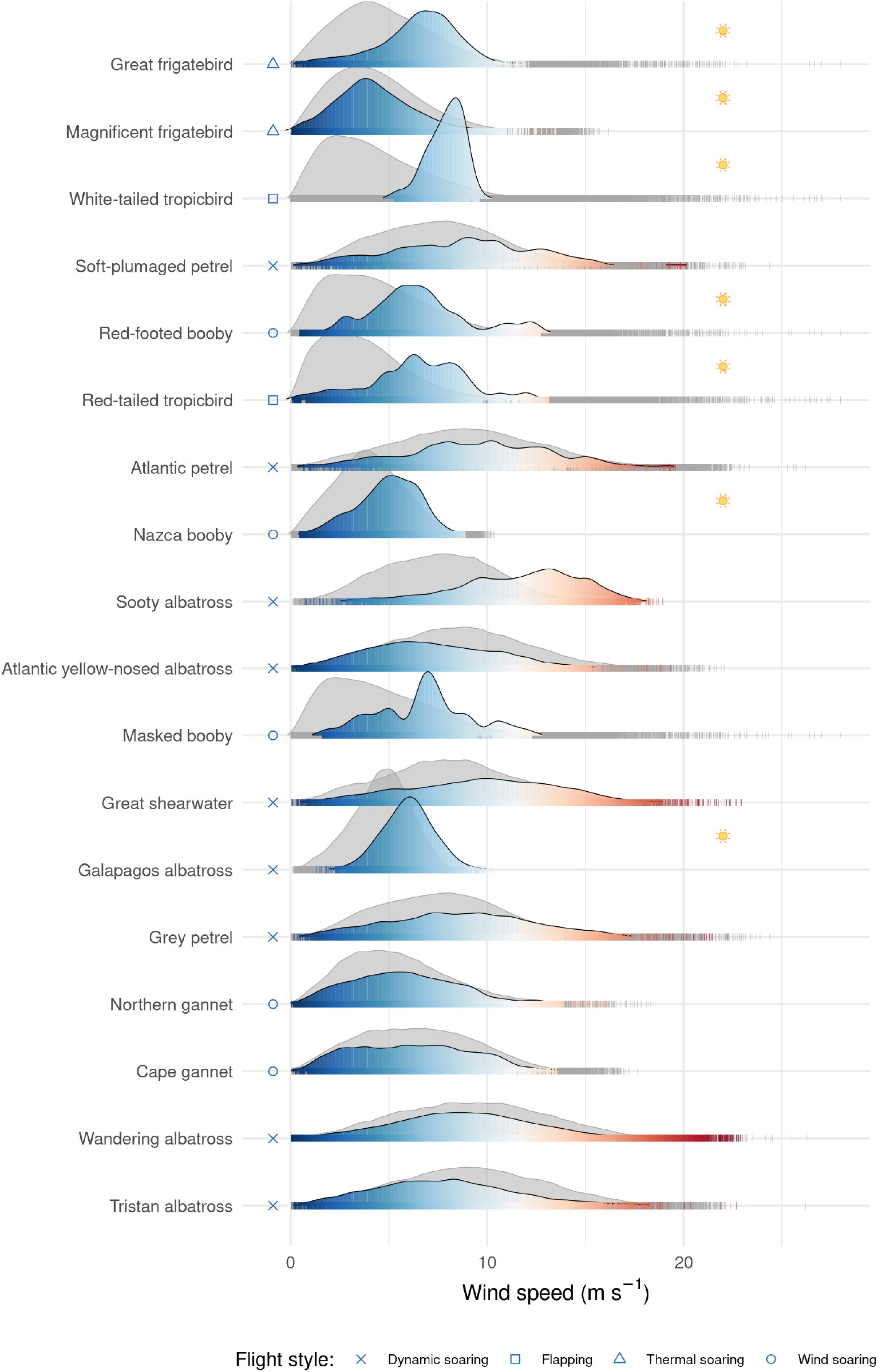
Distribution of wind speeds. Wind speeds experienced during foraging trips (in color. Colors correspond to wind speed, as in Figure 2) is compared to wind conditions across the entire breeding distribution and season over a 5-year period (in gray). Species are ordered by their wing loading, with the lowest wing loading at the top. Tropical species are indicated by a sun icon. See Table S1 for colony locations and wing loadings. See Figure S2 for coefficient of variation of wind speeds encountered by each species.

**Figure 2:**
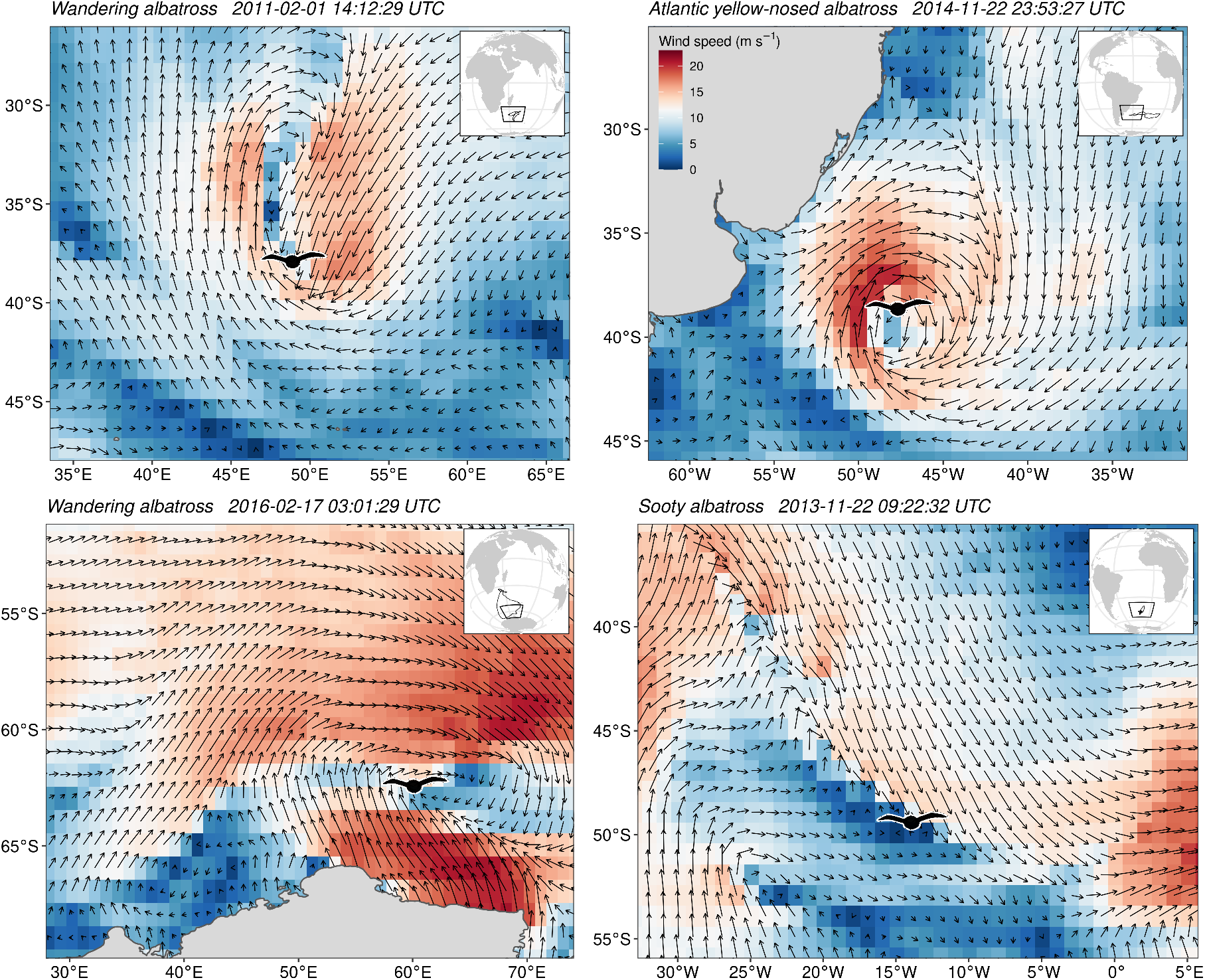
Wind fields at four selected instances where birds avoided the strongest winds. The location of the bird at the time of avoidance of strongest available wind is shown. The full foraging trajectories are shown in the plot insets. In the top two panels birds appear to avoid the strongest winds associated with cyclonic systems by tracking the low wind region in the eye of the storm (see also video S1). In the bottom two panels birds operated along the edge of strong frontal systems, again selecting the region of lower wind speeds. See Figure S3 for available and favored wind speed values.

We then investigated the factors that determined the maximum wind speeds encountered by each species by comparing the encountered maxima with the general wind conditions within the species-specific breeding distributions and seasons. We found that the maximum encountered wind speed was positively correlated with the median (*adj R*^*2*^ = 0.83, p < 0.05), but not the maximum (*adj R*^*2*^ = 0.05, *p* > 0.05), available wind speeds within the breeding range (Table S2). We tested whether morphological characteristics, which largely define flight style, explained species-specific variation in the use of the windscape (Figure 3). Wing loading explained 31% of variation in the strength of tolerable wind speed across species (Table S2). This result was not influenced by temporal and spatial auto-correlation (STAR Methods), or by data quantity (*r* = 0.15, *p* = 0.53).

**Figure 3:**
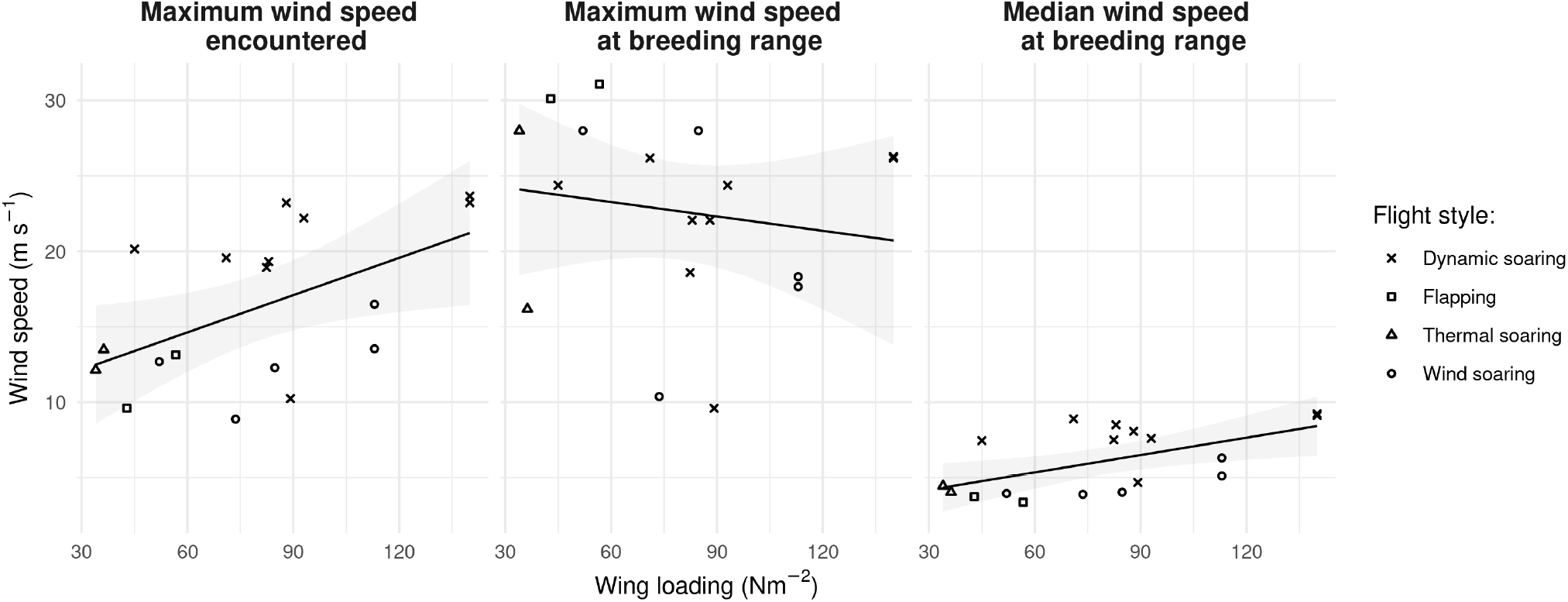
The relationship between wing loading and wind speed. The maximum wind speed encountered by seabirds at the hourly scale was correlated with wing loading (*adj R^2^* = 0.31, *p* < 0.05). Wing loading was predicted by the median (*adj R^2^* = 0.35, *p* < 0.05), but not the maximum (*adj R^2^* = -0.04, *p* = 0.64) wind speeds within the breeding range, indicating that morphological adaptations are a response to the median wind conditions. Shaded areas show the 95% confidence intervals of the regression lines. See Table S2 for complete model summaries.

## Discussion

Against our expectations, we found no general preference or avoidance of specific wind speeds across the 18 species studied at the hourly scale. The lack of a response to the general wind conditions, and the overall low variation in wind speeds available, highlight that the seabird species in our study are adapted to the “wind niche” that they occupy. Indeed, our results suggest that morphology is tightly linked to the wind regime within a breeding distribution, with higher wing loading being advantageous for flight in windier environments. This follows because airspeed increases with wing loading and birds must be able to fly faster than the wind if they are to counter wind drift^13^. Yet while it has been suggested that flight style represents an important aspect of seabird niche space^14^, the role of the windscape in shaping species distributions and diversity has only been investigated for Procellariiformes^15^ (though see ^14^). Our results therefore suggest that wind plays a wider role in the biogeography of flight morphology and flight style through natural selection.

On a day-to-day basis, the strongest winds occurred in the Southern Ocean, an area frequently referred to as the windiest place on earth ^8^. However, while the Southern Ocean had the highest median wind speeds, the overall maximum wind speeds were recorded in the tropical cyclone belt (Figure 1). As a result, the seabirds that are exposed to the strongest global wind strengths (on a seasonal, rather than daily, basis) are those that are morphologically adapted to low median wind speeds, in terms of their ability to counter drift or land safely in strong winds. There should therefore be strong selection pressure for groups such as tropicbirds to develop behavioral adaptations to avoid wind speeds that are some 8 - 9 times the speeds in which they are able to fly. This is consistent with the avoidance of cyclones reported for frigatebirds and boobies, which either remained on land or, depending on their age, made detours of hundreds of kilometers to circumnavigate them^9^.

So why did our step selection approach detect so few instances of wind avoidance? This may be due to the fine scale at which we examined wind speed selectivity (using step lengths of one hour). Given that average airspeeds typically range from 10 to 18 m s^−1^ in seabirds^16^, birds would need to select winds that were significantly different to those available within a radius of some 40 – 65 km, for our approach to categorize the behavior as avoidance. Yet adult Great frigatebirds responded to extreme cyclones (wind speed > 17 m s^−1^) when the storm eye was 250 km away ^9^, and Black-naped terns (*Sterna sumatrana*) departed the colony when cyclones (wind speed > 25 m s^−1^) were 400 to > 2000 km away ^17^. Indeed, it seems likely that the tropical seabirds in this study showed similar long-range responses to extreme winds, or ceased flying altogether, given the discrepancy between the maximum speeds available across the breeding distribution as a whole and those experienced by birds in flight (Figure 1). The use of hourly step lengths may actually select for cases where birds are able to operate very close to storm fronts, which could explain why we found the clearest cases of wind avoidance in albatrosses, the most wind-adapted group.

The few instances where albatrosses did avoid extreme winds provide insight into the speeds and scenarios when wind becomes costly or risky. We show that two species of albatross, including the Wandering albatross, avoided gale force winds of 22 m s^−1^ by flying into and tracking the eye of the storm (Figure 2). Another instance, where an Atlantic yellow-nosed albatross briefly flew into the eye of a weaker storm (around 2014-11-18 21:00, see Video S1), was not identified as wind-avoidance, suggesting that this behavior may be more common than our analyses suggest. Indeed, the same response has been identified in Streaked shearwaters (*Calonectris leucomelas*) ^18^, and is therefore used by species ranging in body mass from 0.5 to 12 kg. Flying towards and remaining within the eye of a storm could be an important part of the behavioral repertoire of fast-flying, wind-adapted species, enabling them to modulate their exposure to unfavorable wind vectors. This raises further interesting questions, such as why birds did not become entrained in the path of the storm. However, understanding how precisely birds interact with the eye of the storm is likely to require wind data at finer temporal resolution than that currently available for public use.

Overall, understanding the responses of animals to global change requires analysis of environmental maxima and rare events. Although temperature maxima and minima are commonly used, e.g. for species distribution modeling ^19^, extreme wind conditions have not received the same attention. While the birds in our study did not select their foraging trajectories in relation to local wind maxima or minima, tropical species did appear to show larger-scale avoidance of extreme winds, as evident in the notable discrepancy between the maximum wind speed they flew in and those available within their breeding distributions. As a result, the maximum wind speeds that birds were recorded flying in provide new insight into the conditions that seabirds with variable size and morphology are able to tolerate. The interspecific variation in these speeds, itself predicted by wing loading, highlights that the definition of “extreme” wind speeds should be considered relative to a given flight morphology, being linked to both flight speed and flight style.

## Acknowledgments

We thank L. Pearmain and B. Clark for assistance with searching and extraction of data from BirdLife International’s Seabird Tracking Database and S. Davidson for assistance with locating relevant Movebank studies. We also thank B. Garde for his input throughout the project. ELCS was supported by a European Research Council starter grant (715874), under the European Union’s Horizon 2020 research and innovation program. EN was supported by the German Academic Exchange Service (DAAD) Postdoctoral Researchers International Mobility Experience (PRIME) fellowship and a Max Planck sabbatical fellowship to ELCS. Data collection from Nazca boobies was supported by the U.S. National Science Foundation under Grant No. DEB 1354473 to DJA. Data collection from Masked boobies was supported by Consejo Nacional de Ciencia y Tecnologia (INAPI-CONACyT) Grant. No. 411876 to ML, the Chilean Millennium Initiative through the Millennium Nucleus Ecology and Sustainable Management of Oceanic Islands (ESMOI), and the Research and Technology Centre (FTZ), University of Kiel. This is a contribution to the Excellence Chair Nouvelle Aquitaine ECOMM led by DG. We thank three anonymous reviewers for their comments and suggestions.

## Author contributions

ELCS conceived the study. EN designed the analyses with input from KS, ELCS, and SdG. DJA, NCC, AF, DG, ML, JLM, LP, PP, NCR, PGR, CDS, SS, VT, HW, and MW contributed data. EN, SdG, and EL prepared the data for analysis. EN analysed the data. EN and ELCS wrote the first draft of the manuscript. All authors commented on and edited the manuscript drafts.

## STAR Methods

### Resource availability

#### Lead contact

Further information and requests for resources and reagents should be directed to and will be fulfilled by the lead contact, Elham Nourani.

#### Materials availability

This study did not generate new unique reagents.

#### Data and code availability

Annotated data are available via the Edmond repository “seabirds’ wind niche” ^20^. R scripts are available via *https://github.com/mahle68/seabirds_storms_public*^21^. Raw datasets are stored on Movebank or the Seabird Tracking Database (as listed in Table S1).

### Experimental model and subject details

All analysis was done on already-existing data collected for free-flying seabirds of the following species:

1. Atlantic yellow-nosed albatross (*Thalassarche chlororhnchosy*)
2. Galapagos albatross (aka. Waved albatross; *Phoebastria irrorata*)
3. Sooty albatross (*Phoebetria fusca*)
4. Tristan albatross (*Diomedea dabbenena*)
5. Wandering albatross (*Diomedea exulans*)
6. Atlantic petrel (*Pterodroma incerta*)
7. Grey petrel (*Procellaria cinerea*)
8. Soft-plumaged petrel (*Pterodroma mollis*)
9. Great shearwater (*Ardenna gravis*)
10. Cape gannet (*Morus capensis*)
11. Northern gannet (*Morus bassanus*)
12. Masked booby (*Sula dactylatra*)
13. Nazca booby (*Sula granti*)
14. Red-footed booby (*Sula sula*)
15. Great frigatebird (*Fregata minor*)
16. Magnificent frigatebird (*Fregata magnificens*)
17. Red-tailed tropicbird (*Phaethon rubricauda*)
18. White-tailed tropicbird (*Phaethon lepturus*)

## Method details

### Bio-logging data

We collected bio-logging data recorded using Platform Transmitter Terminals (PTTs) or GPS loggers for 18 seabird species (Table S1). These datasets were identified by searching Movebank (*www.movebank.org*) and the Seabird Tracking Database (*www.seabirdtracking.org*). The species included in this study represent a range of seabird flight styles (Table S1), including dynamic soaring (gaining energy by repeatedly crossing the vertical gradient of wind velocity over the ocean^22^; nine species of albatross, petrel, and shearwater), wind soaring (regularly alternating flapping and gliding bouts to take advantage of the wind^23^; five species of gannet and booby), thermal soaring (soaring in circles on rising air columns and gliding down in the direction of travel^24^; two species of frigatebird), and obligate flapping fliers (two species of tropicbird). We focused on foraging tracks of adult birds during the breeding season due to the availability of relatively high frequency tracking data, which are scarce outside the breeding season. Diving seabirds such as alcids were not included in our study, as their high wing loading could be selected for due to their use of wings for propulsion underwater and/or to reduce buoyancy^25^.

All data were filtered by speed to ensure that the position information represented periods of flight (threshold of 2 - 3 km h^−1^). To reduce auto-correlation and allow for comparisons between species, we sub-sampled all data to a uniform temporal resolution of at least 1 hour (with tolerance of 15 minutes^26^). In the case of the Galapagos albatross (also known as Waved albatross), we used a 90 minute resolution, which matched the original data frequency. Two species (Red-footed booby and Magnificent frigatebird) were represented from two breeding colonies (Table S1). The Magnificent frigatebird dataset from Isla Contoy had a low temporal resolution (median of 180 minutes) and was excluded from the random step generation procedure (see below). We still report the strongest wind speed encountered by this population and include it in the models for determining exposure to strong wind.

### Processing of bio-logging data

We used a step-selection approach to prepare the data for analysis (Figure S1). This method allowed us to compare the birds’ use of the windscape during foraging trips to conditions that were available, but not used. We considered every two consecutive points along a track as one step, each of them starting at point A and ending at point B. For each of these observed steps, we randomly generated 30 alternative steps, each of which originated in the same place, point A of the observed step, but went to a different location in space, with the same timestamp, as the observed step did. Thus, for each step, we randomly drew 30 values from the distribution of step lengths (Gamma distribution) and turning angles (von Mises distribution) fitted to the empirical data for each species to construct the steps originating in each of the observed location A, but going to 30 alternative B locations. As such, our dataset had a stratified structure, where 30 alternative steps were matched to one observed step per stratum.

For each point in the dataset, we extracted eastward (u) and northward (v) components of wind (at 10 m above surface) from the European Center for Medium-Range Weather Forecast (ECMWF; *www.ecmwf.int*) ERA5 re-analysis database (temporal and spatial resolution of 1 hour and 30 km, respectively). Annotations were done using the ENV-Data track annotation service ^27^ provided by Movebank. We selected bi-linear interpolation for all variables and calculated wind speed using the u and v components of the wind. For each species, we obtained wing loading and aspect ratio from the literature (Table S1).

Preliminary inspection of the largest dataset (Wandering albatross) revealed that the response to extreme winds appeared similar for this species irrespective of whether the step length was set to 1, 2, 4, or 6 hours.

### Wind conditions at species-specific breeding distributions

To understand seabirds’ wind selectivity with regards to the general conditions in their breeding range, we downloaded the eastward and northward components of the wind for the species-specific breeding distributions and seasons. Wind data were obtained for a five year period (2017-2021) from ERA5 for each species breeding range, as given in the BirdLife International database (*http://datazone.birdlife.org*). We then calculated the maximum and median wind speeds within each species’ breeding distribution and season.

### Quantification and statistical analysis

#### Testing for avoidance of strong wind

We used a null modeling technique to test whether seabirds avoided strong winds when foraging at sea. We did not make any assumptions about what wind speeds were considered strong or extreme. Instead, for each set of used and 30 alternative steps (i.e., a stratum), we compared the strongest available wind speed to the wind speed that the individual used. Each stratum was therefore considered to be one sampling unit. Our null hypothesis was that, within each stratum, there was no avoidance of the strongest available wind. Our alternative hypothesis was that the strongest wind available in the stratum was avoided. We calculated a test statistic within each stratum: the difference between the maximum wind speed available and the wind speed at the observed point. To create a null dataset, we grouped the data by species, year, and stratum, and shuffled the wind speed values associated with each row of data within each of these groups. We then calculated the same test statistic within the randomized strata. We repeated the randomization 1,000 times. Given our one-sided alternative hypothesis, we calculated significance as the fraction of random test statistics that were greater than or equal to the observed test statistic.

#### Determinants of exposure to strong wind

We used linear models to investigate whether morphology determined the exposure of seabird species to strong wind conditions. Due to the positive correlation between wing loading and aspect ratio (*r* = 0.60, *p* < 0.05), we included wing loading as the sole predictor in our linear models.

We extracted the maximum wind speed experienced by each species population from the annotated bio-logging dataset (one-hourly resolution). Each population (i.e., species-colony combination) was considered as one sampling unit. We predicted maximum wind as a function of wing loading and checked the model residuals for spatial and temporal auto-correlation that could be related to colony location and timing of breeding (estimated as the median month of the breeding season) following Korner et al.^28^.

The amount of data varied for each species. To test whether the maximum wind speed experienced by different species was affected by the amount of flight data in our dataset, we estimated the correlation between these.

We also investigated whether maximum wind speed experienced by each species was determined by the wind conditions during the breeding season as a whole. For this, we ran linear models predicting maximum encountered wind with the maximum and median wind speed at the breeding distribution.

## Legends for supplemental Excel tables, videos, or dataset files

Video S1: **An Atlantic yellow-nosed albatross flies into the eye of the storm, related to Figure 2**. The bird (indicated by an albatross silhouette) was identified to be avoiding strong winds at 2014-11-22 23:53:27 UTC. However, it also briefly flew into the eye of a weaker storm around 2014- 11-18 21:00 UTC. This instance was not identified as wind-avoidance in our study, suggesting that this behavior may be more common than our analysis suggests. Flying towards and remaining within the eye of a storm could be an important part of the behavioral repertoire of fast-flying, wind-adapted species, enabling them to modulate their exposure to unfavorable winds.

## References

[1] Kawaguchi, K. and Marumo, R. (1964). Mass mortality of Slender-billed Shearwater, Puffinus tenuirostris, in Suruga Bay. J. Yamashina Inst. Ornithol. 4, 106–113.

[2] Ryan, P. and Avery, G. (1987). Wreck of juvenile Blackbrowed Albatrosses on the west-coast of South Africa during storm weather. Ostrich 58, 139–140.

[3] Hume, R. and Christie, D. (1989). Sabine’s Gulls and other seabirds after the October 1987 storm. Br. Birds. 82, 191–208.

[4] Ryan, P., Avery, G., Rose, B., Ross, G.B., Sinclair, J., and Vernon, C. (1989). The Southern Ocean seabird irruption to South African waters during winter 1984. Mar. Ornithol. 17, 41–55.

[5] Camphuysen, C., Wright, P., Leopold, M., Hüppop, O., and Reid, J. (1999). A review of the causes, and consequences at the population level, of mass mortalities of seabirds. Ices coop res report no 232 edition (International Council for the Exploration of the Sea, Copenhagen).

[6] Hass, T., Hyman, J., and Semmens, B. (2012). Climate change, heightened hurricane activity, and extinction risk for an endangered tropical seabird, the black-capped petrel Pterodroma hasitata. Mar. Ecol. Prog. Ser. 454, 251–261. doi: 10.3354/meps09723.

[7] Young, I.R. and Ribal, A. (2019). Multiplatform evaluation of global trends in wind speed and wave height. Science 364, 548–552. doi: 10.1126/science.aav9527.

[8] Weimerskirch, H., Guionnet, T., Martin, J., Shaffer, S.A., and Costa, D. (2000). Fast and fuel efficient? Optimal use of wind by flying albatrosses. Proc. Royal Soc. B. 267, 1869–1874. doi: 10.1098/rspb.2000.1223.

[9] Weimerskirch, H. and Prudor, A. (2019). Cyclone avoidance behaviour by foraging seabirds. Sci. Rep. 9, 1–9. doi: 10.1038/s41598-019-41481-x.

[10] Tarroux, A., Weimerskirch, H., Wang, S.H., Bromwich, D.H., Cherel, Y., Kato, A., Ropert-Coudert, Y., Varpe, Ø., Yoccoz, N.G., and Descamps, S. (2016). Flexible flight response to challenging wind conditions in a commuting Antarctic seabird: do you catch the drift? Anim. Behav. 113, 99–112. doi: 10.1016/j.anbehav.2015.12.021.

[11] Amélineau, F., Péron, C., Lescroël, A., Authier, M., Provost, P., and Grémillet, D. (2014). Windscape and tortuosity shape the flight costs of northern gannets. J. Exp. Biol. 217, 876–885. doi: 10.1242/jeb.097915.

[12] Thurfjell, H., Ciuti, S., and Boyce, M.S. (2014). Applications of step-selection functions in ecology and conservation. Mov. Ecol. 2, 1–12. doi: 10.1186/2051-3933-2-4.

[13] Tennekes, H. (2009). The Simple Science of Flight, Revised and Expanded Edition: From Insects to Jumbo Jets (Cambridge: MIT press).

[14] Spear, L.B. and Ainley, D.G. (1998). Morphological differences relative to ecological segregation in petrels (Family: Procellariidae) of the Southern Ocean and tropical Pacific. Auk 115, 1017–1033.

[15] Davies, R.G., Irlich, U.M., Chown, S.L., and Gaston, K.J. (2010). Ambient, productive and wind energy, and ocean extent predict global species richness of Procellariiform seabirds. Glob. Ecol. Biogeogr. 19, 98–110. doi: 10.1111/j.1466-8238.2009.00498.x.

[16] Spear, L.B. and Ainley, D.G. (1997). Flight speed of seabirds in relation to wind speed and direction. Ibis 139, 234–251. doi: 10.1111/j.1474-919X.1997.tb04621.x.

[17] Thiebot, J.B., Nakamura, N., Toguchi, Y., Tomita, N., and Ozaki, K. (2020). Migration of black-naped terns in contrasted cyclonic conditions. Mar. Biol. 167, 1–12. doi: 10.1007/s00227-020-03691-0.

[18] Lempidakis, E., Shepard, E.L.C., Ross, A.N., Matsumoto, S., Koyama, S., Takeuchi, I., and Yoda, K. (2022). Pelagic seabirds reduce risk by flying into the eye of the storm. PNAS 119, e2212925119. doi: 10.1073/pnas.2212925119.

[19] Fick, S.E. and Hijmans, R.J. (2017). WorldClim 2: new 1-km spatial resolution climate surfaces for global land areas. Int. J. Climatol. 37, 4302–4315. doi: 10.1002/joc.5086.

[20] Nourani, E. (2022). seabirds’ wind niche. https://doi.org/10.17617/3.8U7EHD.

[21] Nourani, E. (2023). mahle68/seabirds_storms_public: Seabirds’ wind niche (v1.0). Zenodo. https://doi.org/10.5281/zenodo.7538390

[22] Richardson, P.L., Wakefield, E.D., and Phillips, R.A. (2018). Flight speed and performance of the wandering albatross with respect to wind. Mov. Ecol. 6, 1–15. doi: 10.1186/s40462-018-0121-9.

[23] Weimerskirch, H., Le Corre, M., Ropert-Coudert, Y., Kato, A., and Marsac, F. (2005). The three-dimensional flight of red-footed boobies: adaptations to foraging in a tropical environment? Proc. Royal Soc. B. 272, 53–61. doi: 10.1098/rspb.2004.2918.

[24] Weimerskirch, H., Chastel, O., Barbraud, C., and Tostain, O. (2003). Frigatebirds ride high on thermals. Nature 421, 333–334. doi: 10.1038/421333a.

[25] Lapsansky, A.B., Warrick, D.R., and Tobalske, B.W. (2022). High Wing-Loading Correlates with Dive Performance in Birds, Suggesting a Strategy to Reduce Buoyancy. Integr. Comp. Biol. 62, 878–889. doi: 10.1093/icb/icac117.

[26] Kranstauber, B., Smolla, M., and Scharf, A.K. (2020). Package ‘move’.

[27] Dodge, S., Bohrer, G., Weinzierl, R., Davidson, S.C., Kays, R., Douglas, D., Cruz, S., Han, J., Brandes, D., and Wikelski, M. (2013). The environmental-data automated track annotation (Env-DATA) system: linking animal tracks with environmental data. Mov. Ecol. 1, 3. doi: 10.1186/2051-3933-1-3.

[28] Korner-Nievergelt, F., Roth, T., Von Felten, S., Guélat, J., Almasi, B., and Korner-Nievergelt, P. (2015). Bayesian data analysis in ecology using linear models with R, BUGS, and Stan (Cambridge: Academic Press).

