## supplementary_material for "Seabird morphology determines operational wind speeds, tolerable maxima and responses to extremes"

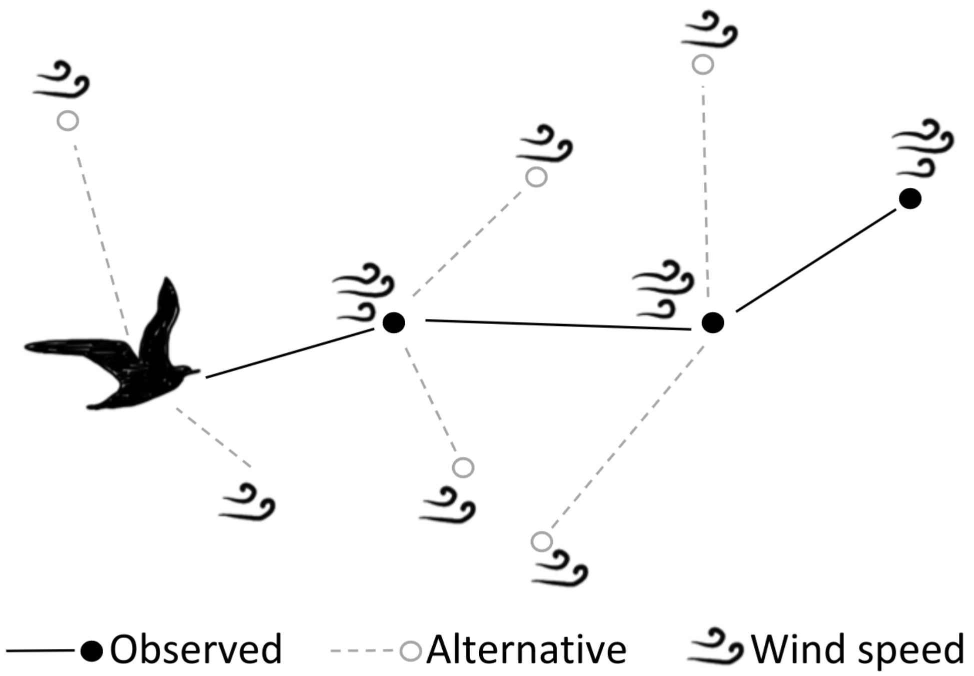


Figure S1: S**chematic example of the step-selection approach used in this study, related to the STAR Methods**. Each pair of consecutive locations is considered as one observed step. Each observed step is associated with a number of alternative steps (2 in this example), creating one stratum. The location of alternative steps is determined by randomly selecting a step length and a turn angle from the gamma distribution of step lengths and the von Mises distribution of turn angles observed for each species. The favored wind speed within each stratum is then compared to the maximum available wind speed within the stratum. In this example the bird favored the strongest available wind in all the strata.

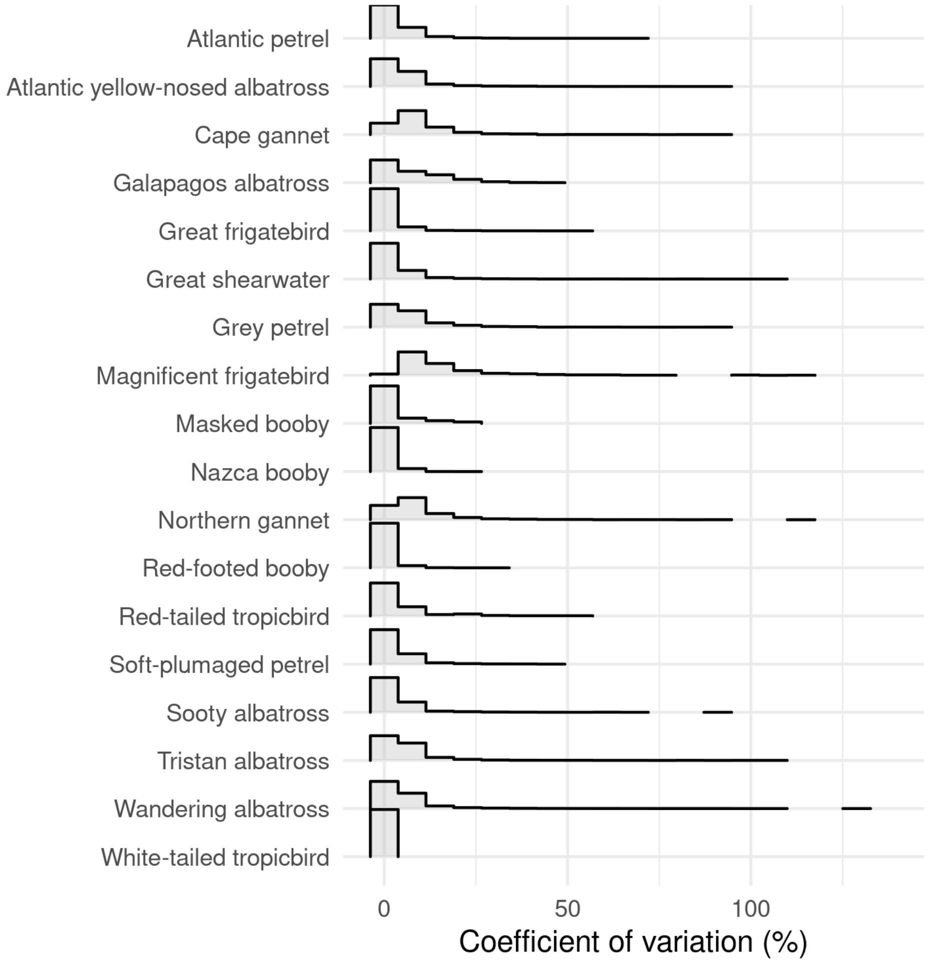


Figure S2: **Distribution of coefficient of variation estimated for wind speed within each stratum for each species, Related to Figure 1.** Values exceeding 100% indicate higher standard deviation than the mean. See Figure S2 for results of null modeling to set for avoidance of strong winds within the strata.

**(A)**

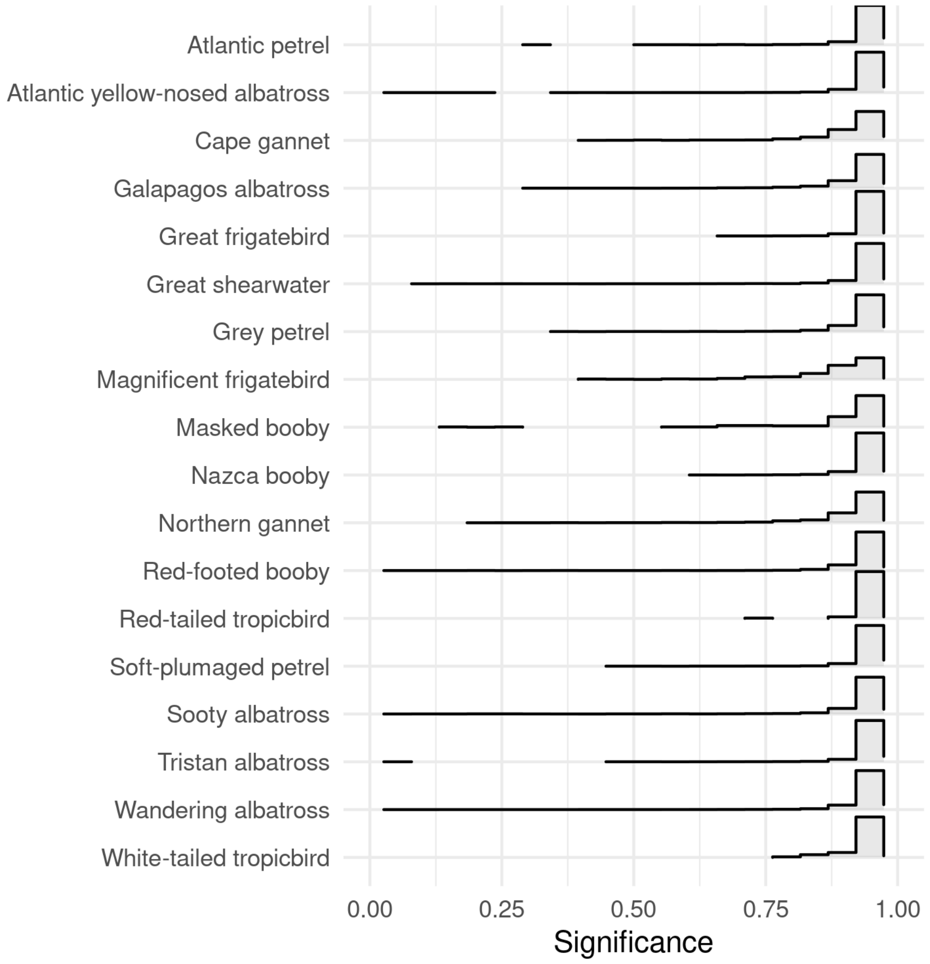


**(B)**

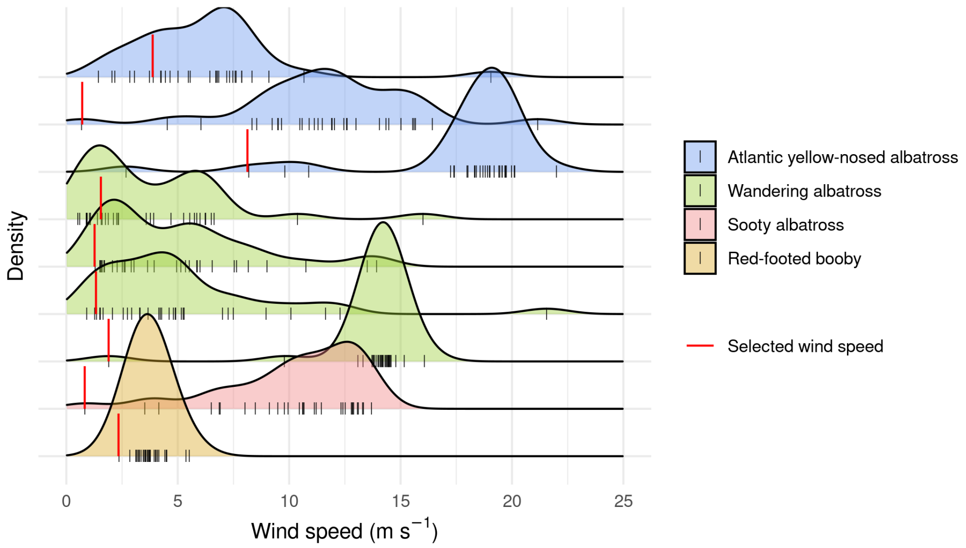


Figure S3: **Results of the randomization test, related to Figure 2.** (A) Significance values for randomization tests performed on the stratified dataset to test for avoidance of strong wind The null hypothesis was accepted in the majority of strata: seabirds did not avoid strong winds. In fact, significant values of 1 indicate that the bird used the maximum wind speed available. (B) Distribution of available wind speed at the strata where strong wind was avoided. Wandering albatrosses sometimes avoided wind speeds that were well within their normal operational range (14, 15, and 16 m $s^{-1}$). This could either be because individual trajectories were driven by factors other than the wind field in these instances, or because birds chose to select/avoid wind conditions based on the direction rather than speed alone, for instance to enable the efficient exploitation of cross winds.

Table S1. **Details of the data used in the study, relates to the STAR Methods.** Database names, sources, and additional information are provided as footnotes. References are indicated by numbers proceeding an "S".

| Species | Colony | Device | Frequency (median; mins) | Years | Individuals (n) | Tracks (n) | Wing loading ($Nm^{-2}$) | Aspect ratio | Flight style |
|---|---|---|---|---|---|---|---|---|---|
| Atlantic yellow-nosed albatross | Gough Island | GPS | 60 | 2013-2014 | 39 | 49 | 83.00 [S1] | 15.1 [S1] | dynamic soaring |
| Galapagos albatross [c] [S2] | Isla Espanola | GPS | 90 | 2008 | 13 | 91 | 89.20 [S3] | 14.7 [S3] | dynamic soaring |
| Sooty albatross | Gough Island | GPS | 11 | 2013 | 13 | 17 | 82.40 [f] | 14.1 [f] | dynamic soaring |
| Tristan albatross | Gough Island [a] | GPS | 60 | 2014 | 18 | 48 | 140.00 [b] | 15 [b] | dynamic soaring |
| Wandering albatross | Crozet Island | GPS | 2 | 2010-2019 | 417 | 471 | 140.00 [S4] | 15 [S4] | dynamic soaring |
| Atlantic petrel | Gough Island | GPS | 60 | 2014 | 5 | 5 | 71.00 [m] | 11.4 [m] | dynamic soaring |
| Grey petrel | Gough Island | GPS | 30 | 2014 | 15 | 15 | 93.00 [S1] | 12.1 [S1] | dynamic soaring |
| Soft-plumaged petrel | Gough Island | GPS | 60 | 2014 | 8 | 9 | 45.00 [S1] | 10.5 [S1] | dynamic soaring |
| Great shearwater | Gough Island | GPS & PTT | 60 | 2009-2010, 2013-2014 | 26 | 31 | 88.00 [k] | 11 [S5] | dynamic soaring |
| Cape gannet [d] | Algoa Bay | GPS | 0.02 | 2005& 2009 | 56 | 56 | 113.00 [e] | 13.1 [e] | wind soaring |
| Northern gannet [d] | Ile Rouzic | GPS | 0.5 | 2005-2017 | 150 | 150 | 113.00 [S6] | 13.1 [S6] | wind soaring |
| Masked booby [g] [S7] | Motu Nui | GPS | 4 | 2016-2017 | 23 | 98 | 84.76 [S8] | 12.2 [S8] | wind soaring |
| Nazca booby [h] | Isla Espanola | GPS | 3 | 2011-2017 | 55 | 122 | 73.64 [S9] | 7.10 [S9] | wind soaring |
| Red-footed booby | Europa Island; Genovesa Island | GPS | 0.2; 2 | 2009, 2012, 2014 | 95 | 126 | 52.00 [S6] | 11.3 [S6] | wind soaring |
| Great frigatebird [j] [S10] | Genovesa Island | PTT | 54 | 2015-2016 | 11 | 47 | 34.00 [S6] | 11.1 [S6] | thermal soaring |
| Magnificent frigatebird [i] | Iguana Island; Isla Contoy | GPS | 5; 180 | 2011-2016 | 14 | 234 | 36.28 [S11] | 12.0 [S11] | thermal soaring |
| Red-tailed tropicbird [l] | Round Island | GPS | 1 | 2018 | 55 | 76 | 56.67 [S8] | 10.9 [S8] | flapping |
| White-tailed tropicbird | Fernando de Noronha | GPS | 1 | 2015 | 15 | 18 | 42.86 [S12] | 10.1 [S12] | flapping[l] |

[a] All data from Gough Island are collected by Peter Ryan; available via *http://www.seabirdtracking.org/*

[b] Data for Wandering albatross was used [S4].

[c] Movebank Study: "Galapagos Albatrosses"

[d] Data collected by David Gremillet; available via *http://www.seabirdtracking.org/*

[e] Data for Northern gannet was used [S6].

[f] Data for light-mantled sooty albatross was used [S4].

[g] Movebank Study: "Foraging ecology of masked boobies (Sula dactylatra) in the world's largest “oceanic desert”"

[h] Movebank Study: "Nazca booby Sula granti Isla Espanola, Galapagos."

[i] Movebank Studies: "MPIAB PNIC hurricane frigate tracking" and "Frigatebirds breeding at Iguana Island, Panama"

[j] Movebank Study: "Great frigatebirds (Weimerskirch)"

[k] Data for sooty shearwater [S6].

[l] Movebank Study: "Red-tailed tropicbirds (Phaethon rubricauda) Round Island"

[m] Data for White-headed petrel was used [S1].

[l] White-tailed tropicbirds have been found to undertake some thermal soaring.

Table S2. **Summary of the linear models, related to Figure 3.** Max wind encountered refers to the maximum wind speed that the bird encountered during foraging trips (based on bio-logging data). Median and Maximum wind at breeding range were estimated based on 5 years of wind data downloaded for the species-specific breeding distribution range during the breeding season.

| Dependent variable | Predictor variable | $\beta$ (Std. Error) | $R^2$ | Adjusted $R^2$ | Residual Std. Error | F Statistic |
|---|---|---|---|---|---|---|
| Max wind encountered | Wing loading | $0.34^{**}$ (0.11) | 0.35 | 0.31 | 15.57 (df = 18) | $9.54^{***}$ (df = 1;18) |
| Wing loading | Median wind at breeding | $9.40^{**}$ (2.82) | 0.38 | 0.35 | 26.39 (df = 18) | $11.06^{***}$ (df = 1;18) |
| Wing loading | Maximum wind at breeding | -0.58 (1.22) | 0.01 | -0.04 | 33.32 (df = 18) | 0.23 (df = 1;18) |
| Max wind encountered | Median wind at breeding | $8.01^{***}$ (0.82) | 0.84 | 0.83 | 7.69 (df = 18) | $94.91^{***}$ (df = 1;18) |
| Max wind encountered | Maximum wind at breeding | 0.65 (0.69) | 0.05 | 0 | 18.79 (df = 18) | 0.90 (df = 1;18) |
| *Note:* | $^{*}$p<0.1; $^{**}$p<0.05; $^{***}$p<0.01 | | | | | |

### References


[S1] Warham, J. (1977). Wing loadings, wing shapes, and flight capabilities of Procellariiformes. New Zealand Journal of Zoology *4*, 73–83.

[S2] Cruz, S., Proaño, C., Anderson, D., Huyvaert, K., and Wikelski, M. (2013). Data from: The Environmental-Data Automated Track Annotation (Env-DATA) System: Linking animal tracks with environmental data. doi `doi:10.5441/001/1.3hp3s250`.

[S3] Suryan, R.M., Anderson, D.J., Shaffer, S.A., Roby, D.D., Tremblay, Y., Costa, D.P., Sievert, P.R., Sato, F., Ozaki, K., Balogh, G.R., et al. (2008). Wind, waves, and wing loading: morphological specialization may limit range expansion of endangered albatrosses. PLoS One *3*, e4016. doi `10.1371/journal.pone.0004016`.

[S4] Pennycuick, C.J. (1982). The flight of petrels and albatrosses (Procellariiformes), observed in South Georgia and its vicinity. Philosophical Transactions of the Royal Society of London. B, Biological Sciences *300*, 75–106.

[S5] Alerstam, T., Gudmundsson, G.A., and Larsson, B. (1993). Flight tracks and speeds of Antarctic and Atlantic seabirds: radar and optical measurements. Philosophical Transactions of the Royal Society of London. Series B: Biological Sciences *340*, 55–67.

[S6] Spear, L.B. and Ainley, D.G. (1997). Flight behaviour of seabirds in relation to wind direction and wing morphology. Ibis *139*, 221–233.

[S7] Lerma, M., Serratosa, J., Luna-Jorquera, G., and Garthe, S. (2020). Foraging ecology of masked boobies (Sula dactylatra) in the world's largest "oceanic desert". Marine Biology *167*, 1–13. doi `10.1007/s00227-020-03700-2`.

[S8] Hertel, F. and Ballance, L.T. (1999). Wing ecomorphology of seabirds from Johnston Atoll. The Condor *101*, 549–556.

[S9] Howard, J.L., Tompkins, E.M., and Anderson, D.J. (2021). Effects of age, sex, and ENSO phase on foraging and flight performance in Nazca boobies. Ecology and evolution *11*, 4084–4100. doi 10.1002/ece3.7308.

[S10] Weimerskirch, H., Borsa, P., Cruz, S., de Grissac, S., Gardes, L., Lallemand, J., Corre, M.L., and Prudor, A. (2017). Diversity of migration strategies among great frigatebirds populations. Journal of Avian Biology *48*, 103–113. doi 10.1111/jav.01330.

[S11] Trefry, S.A. and Diamond, A.W. (2017). Exploring hypotheses for sexual size dimorphism in frigatebirds. Evolutionary Ecology Research *18*, 225–252.

[S12] Pennycuick, C.J. (2008). Modelling the flying bird (London: Academic Press).
